# Selective octopaminergic tuning of mushroom body circuits during memory formation

**DOI:** 10.1101/2025.06.16.659845

**Authors:** Ulrike S. Franke, Alexandra Großjohann, Samantha Aurich, Ines Köhler, Marius Lamberty, Mareike Selcho, Robert J. Kittel, Andreas S. Thum, Dennis Pauls

**Affiliations:** Department of Animal Physiology, Institute of Biology, Leipzig University, Leipzig, Germany; Department of Genetics, Institute of Biology, Leipzig University, Leipzig, Germany; German Centre for Integrative Biodiversity Research (iDiv) Halle-Jena-Leipzig, Leipzig, Germany

## Abstract

The catecholamines octopamine and tyramine undoubtedly have a major impact on the life of an insect. A wide range of physiological processes and behaviours are regulated by these neurotransmitters/hormones. Octopamine and tyramine act homologous to the adrenergic system of vertebrates, primarily adapting the organism to the given situation, by switching between the states of alertness and rest. Interestingly, higher brain functions like learning and memory are also regulated by octopamine and tyramine. About 30 years ago, initial work in *Drosophila* has demonstrated that dopaminergic neurons signal punishment, while octopaminergic neurons signal reward during olfactory associative learning and memory. In the meantime, however, it has become clear that distinct types of dopaminergic neurons convey both reward and punishment signals to the mushroom bodies, a central brain region responsible for the formation and storage of associative memories. Although some conflicting data remain, these findings challenge the previously established model of functional segregation and may limit the proposed role of octopamine neurons as teaching neurons during memory formation. We have therefore re-examined the role of octopamine in learning and memory in *Drosophila* larvae. Through a combination of Ca^2+^ imaging, anatomical studies and gain-of-function and loss-of-function behavioural approaches, we demonstrate that octopamine signalling plays a crucial role in larval learning by modulating dopaminergic neurons across distinct cell clusters to orchestrate memory processes.

## INTRODUCTION

On the basis of several key studies, it has become an accepted principle that memory formation in *Drosophila* is regulated by the differential involvement of distinct catecholamines. This has been demonstrated in both *Drosophila* larvae (Honjo and Furukubo-Tokunaga, 2009; Schroll et al., 2006) and flies (reviewed in (Heisenberg, 2003; Waddell, 2013); (Schwaerzel et al., 2003)) as well as in other invertebrate species (e.g. (Agarwal et al., 2011; Hammer, 1993; Menzel, 2012; Mizunami et al., 2009; Unoki et al., 2006, 2005; Vergoz et al., 2007; Wissink and Nehring, 2021; Yu et al., 2022)). Remarkably, the proposed functional separation for octopamine (OA; mediating reward during appetitive learning) and dopamine (DA; mediating punishment during aversive learning) did not withstand further testing. Several studies have convincingly shown that DA also plays a critical role in punishment and reward learning in both *Drosophila* larvae and flies (Burke et al., 2012; Huetteroth et al., 2015; Ichinose et al., 2015; Kim et al., 2007; Liu et al., 2012; Lyutova et al., 2019; Rohwedder et al., 2016; Selcho et al., 2009; Yamagata et al., 2015). Specifically, larvae and flies lacking the dopamine type-1 receptor dDA1 display a reduced performance not only in punishment learning but also in reward learning (Kim et al., 2007; Selcho et al., 2009). In addition, selectively silencing DA neurons of the protocerebral anterior medial (PAM) cluster only during training specifically impaired appetitive learning in both larvae and flies (Burke et al., 2012; Ichinose et al., 2015; Liu et al., 2012; Rohwedder et al., 2016; Saumweber et al., 2018). Overall, these results suggest a developmentally conserved wiring principle in the *Drosophila* brain, where two distinct clusters of DA neurons in the *Drosophila* brain mediate appetitive and aversive reinforcement: the PPL1 (protocerebral posterior lateral 1) and PAM clusters in adults, and the DL1 (dorsolateral 1) and primary PAM (pPAM) clusters in larvae.

Yet, what is the role of OA, and how should the previous findings be interpreted in this context? The findings of Burke and colleagues suggest that the OA-dependent effect is upstream of DA neurons, as memory formation induced by thermogenetic activation of OA neurons was impaired in a dopamine receptor (dDA1/*dumb^1^*) mutant background (Burke et al., 2012). Furthermore, blockage or ablation of OA neurons disrupted appetitive learning in larvae and flies when arabinose - a sweet-tasting sugar with no nutritional value - was used as a reward (Burke et al., 2012; Selcho et al., 2014). In contrast, appetitive learning with nutritive sugars such as sucrose or fructose remained unaffected. Interestingly, odour-arabinose learning was impaired following the downregulation of the OA receptor OAMB in PAM neurons, indicating that OAMB is required in DA neurons for the OA-dependent memory formation (Burke et al., 2012). Collectively, these findings suggest that the sweetness of a sugar is conveyed via OA neurons to the PAM cluster through OAMB, where it is integrated with additional sensory information to generate a positive reinforcement signal. This model was further refined by Huetteroth and colleagues, who demonstrated that the OA-dependent sweetness signal specifically establishes short-term memory via OAMB-mediated signaling in a subset of PAM neurons that innervate the ß’_2am_ and y_4_ regions of the mushroom body (Huetteroth et al., 2015). However, another OA receptor, Octß2R, also appears to be involved. Knockdown of Octβ2R in MB-MP1-type DA neurons similarly impaired reward learning (Burke et al., 2012). The dopaminergic MB-MP1 neurons not only contribute to odour-specific aversive memory formation but also relay information about the animaĺs nutritional state to the mushroom body as they are modulated by dNPF (*Drosophila* neuropeptide F), a neuropeptide representing hunger state. These findings imply that OA signaling might encode more than a simple positive teaching signal via OAMB; it may also modulate negative reinforcement or hunger-related signals via Octβ2R. Nonetheless, this model requires further refinement based on several additional findings. First, *oamb* mutant flies show severely impaired sucrose learning, and this deficit can be rescued by expressing OAMB in intrinsic mushroom body Kenyon cells, suggesting an OA-dependent function that operates independently of DA neurons (Kim et al., 2013). On top, OA is also essential for aversive olfactory learning. Flies lacking tyramine-ß-hydroxylase (TßH), the enzyme hydroxylating tyramine (TA) to OA, exhibit impaired odour-shock memory, which could be rescued by targeted TßH expression in OA neurons (Iliadi et al., 2017). Moreover, *Octβ1R* mutant flies display deficits in aversive odor-shock memory, and these impairments could be rescued by *Octβ1R* expression in Kenyon cells (Sabandal et al., 2020). Finally, the anterior paired lateral (APL) neuron, which innervates the whole mushroom body, was shown to be octopamine-positive and function via Octß2R in Kenyon cells in anesthesia-resistant memory in flies (Wu et al., 2013), providing further evidence for an OA-dependent mechanism that functions independently of DA neurons.

Given the diverse modes of action attributed to OA, we focused this study on re-examining the role of OA in *Drosophila* larval learning. To this end, we employed a combination of optogenetics, loss-of-function behavioural assays, and Ca^2+^ imaging. Our results indicate that OA neurons do not act as direct teaching neurons during memory formation. Instead, OA modulates distinct clusters of DA neurons, thereby adding an additional layer of complexity to the mushroom body circuitry underlying learning and memory. Consequently, the function of OA in learning appears to centre on the contextual adaptation of neuronal processes, such as aligning behavioural responses with internal states and physiological needs.

## RESULTS

Previous work on larval learning has comprehensively shown that associative olfactory memories are stored within the mushroom bodies (Eichler et al., 2017; Pauls et al., 2010b; Widmann et al., 2018). Moreover, DA neurons of the DL1 cluster provide an aversive teaching signal (Honjo and Furukubo-Tokunaga, 2009; Selcho et al., 2009; Weber et al., 2023b) while DA neurons of the pPAM cluster convey an appetitive one (Lyutova et al., 2019; Rohwedder et al., 2016; Saumweber et al., 2018; Schleyer et al., 2020). However, this working hypothesis somehow neglects the role of OA, despite clear evidence that OA signalling is essential for establishing larval reward memories (Honjo and Furukubo-Tokunaga, 2009; Selcho et al., 2014). A conundrum arises from anatomical studies: neither light microscopy nor EM data provide evidence for direct innervation of the medial lobes by OA neurons. In contrast, the medial lobes - mushroom body compartments where reward memories are stored - are directly innervated by pPAM neurons (Eichler et al., 2017; Selcho et al., 2014). Nevertheless, the prevailing mediation hypothesis posits that OA neurons convey teaching signals directly to mushroom body Kenyon cells during memory formation.

To initially set aside any specific role of OA neurons during memory formation and instead confirm their general involvement, we optogenetically activated Tdc2-positive neurons **(Fig.S1A-H)** during training. Tyrosine decarboxylase (Tdc) is crucial for the synthesis of TA, thus *Tdc2-GAL4* includes all TA and OA neurons (Cole et al., 2005; Selcho et al., 2014, 2012). In detail, we employed a well-established reciprocal two-group design **(Fig.1A),** in which one odour was coupled with blue light exposure (CS+) and the other was presented under dim red light (CS-) (Rohwedder et al., 2016; Schroll et al., 2006). Following the protocol by Lyutova et al. (Lyutova et al., 2019), we applied blue light (475nm) at an intensity of ∼10^-1^mW/mm^2^ to induce neuronal activity through the activation of the channelrhodopsin XXL variant (chop^XXL^; (Dawydow et al., 2014)). CantonS wildtype larvae showed no significant approach or avoidance, suggesting that light exposure at this intensity is not sufficient to elicit memory expression (Lyutova et al., 2019; von Essen et al., 2011) **(Fig.1B)**. Surprisingly, and in contrast to previously published data (Schroll et al., 2006), *Tdc2>chop^XXL^*larvae did not show any memory expression in our substitution learning assay **(Fig.1C)**. As a positive control, we used optogenetic activation of DA neurons from the pPAM cluster (*R58e02>chop^XXL^*; **Fig.S1I-P**), which have been shown to mediate reward signals to the medial lobe of the mushroom bodies (Lyutova et al., 2019; Rohwedder et al., 2016; Saumweber et al., 2018). In line with previous reports, *R58e02>chop^XXL^* larvae showed robust appetitive memory expression **(Fig.1C)**.

**Figure 1:**
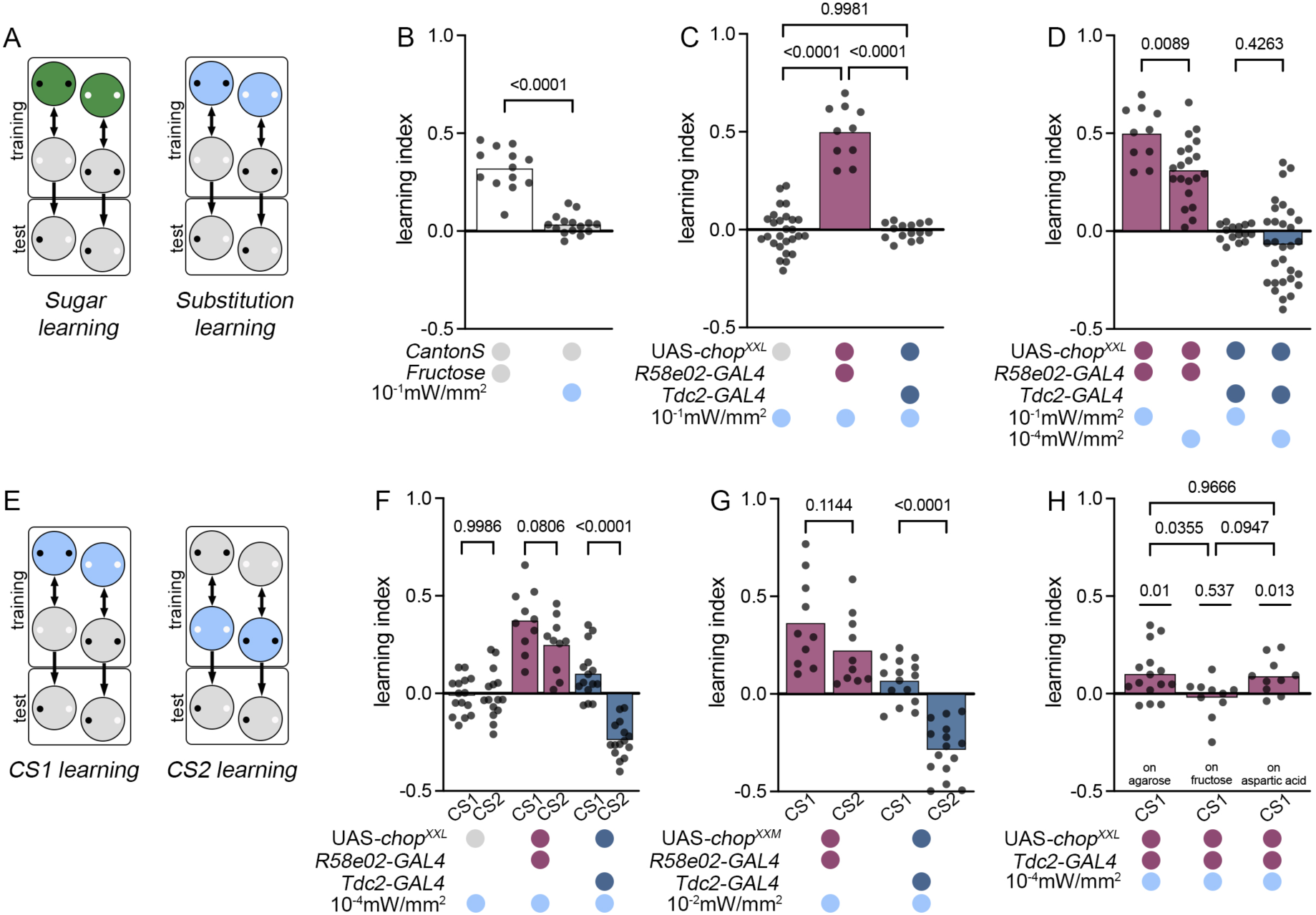
Optogenetic activation of OA neurons induces memory expression. (A) Schematic overview of learning protocols used in this study. (B) Wildtype larvae formed and expressed odour-sugar memories (white plot), while no memory expression was detectable with coupled odour and blue light (grey plot; 10^-1^mW/mm^2^, 475nm). (C) Blue light exposure substituted for sugar presentation in R58e02>chop^XXL^ larvae due to the expression of channelrhodopsin in PAM neurons. In contrast, optogenetic activation of octopaminergic neurons was not sufficient to induce memory expression in Tdc2>chop^XXL^ larvae. (D) R58e02>chop^XXL^ larvae showed memory expression with lower light intensity (10^-4^mW/mm^2^), but significantly reduced compared to 10^-1^mW/mm^2^ of blue light. Tdc2>chop^XXL^ larvae showed no memory expression. (E) Schematic illustration of CS1 and CS2 learning. During CS1 learning, the first presented odour is coupled to the US, whereas during CS2 learning, the second presented odour is coupled to the US. (F) While pairing blue light activation with the first odour (CS1 learning) elicited reward memory expression, pairing the second presented odour elicited aversive memory expression in Tdc2>chop^XXL^ larvae. No sequence effect was detectable for dopaminergic neurons in R58e02>chop^XXL^ larvae. (G) Similar results were obtained using chop^XXM^ indicating that the sequence effect is not based on the long open state of channelrhodopsin XXL. (H) The optogenetically induced appetitive CS1 memory is expressed on a pure agarose and aspartic acid test plate, whereas no memory expression is seen on a fructose test plate.

Although high-intensity blue light has been successfully used to substitute the US during learning by activating mushroom body Kenyon cells (Lyutova et al., 2019), *chop^XXL^* has been shown to function more efficiently at light intensities three orders of magnitude lower than previous variants, such as chop-wt (Dawydow et al., 2014). Furthermore, previous studies have demonstrated that both light intensity and frequency are critical factors influencing the strength of the behavioural outcome following the optogenetic activation of OA neurons (Claßen and Scholz, 2018). We therefore repeated the substitution learning, but this time with reduced light intensity **(Fig.1D)**. *R58e02>chop^XXL^* exhibited appetitive memory expression using blue light with an intensity of ∼10^-4^mW/mm^2^. Interestingly, learning scores were significantly lower compared to those obtained with an intensity of 10^-1^mW/mm^2^. In contrast, no memory expression was detected in *Tdc2>chop^XXL^* larvae at the lower light intensity **(Fig.1D)**. However, we noted that data variability in *Tdc2>chop^XXL^* was different between the two light intensities. Our previous work has shown that learning performance can vary depending on whether the US is paired with the first odour (CS1 learning; **Fig.1E**) or the second odour (CS2 learning; **Fig.1E**) during training (Pauls et al., 2010a). Therefore, we tested whether sequence effects could account for the increased data variability observed with 10^-4^mW/mm^2^ light intensity. Indeed, *Tdc2>chop^XXL^* larvae showed significantly different performance depending on whether the 1^st^ odour was paired with blue light (CS1 learning) or the 2^nd^ odour (CS2 learning; **Fig.1E-F**). Surprisingly, CS1 learning resulted in a positive learning score, whereas CS2 learning induced a negative learning score **(Fig.1F),** suggesting that OA neurons can induce both appetitive and aversive memory formation, depending on the timing of activation. In contrast, no sequence effect was detected in *R58e02>chop^XXL^*larvae. To further validate the OA neuron-specific effect, we used chop^XXM^ to optogenetically activate OA neurons (Scholz et al., 2017). Consistently, activation of Tdc2-positive neurons with *chop^XXM^* (with ∼10^-2^mM/mm^2^ of blue light) induced appetitive memory expression for CS1 learning and aversive memory expression for CS2 learning **(Fig.1G)**. These findings suggest that the observed sequence effect is not due to the prolonged open state or extra high expression levels of *chop^XXL^* (Dawydow et al., 2014). Next, we tested the specificity of the optogenetically induced CS1 memory in *Tdc2>chop^XXL^* larvae by presenting fructose or aspartic acid in the test situation. While the presence of fructose impaired memory expression, aspartic acid had no effect, indicating that the memory is specifically related to reward quality and valence (Schleyer et al., 2015) **(Fig.1H)**. Taken together, these findings suggest that OA neurons are involved in both larval appetitive and aversive memory formation.

Using single cell anatomical description of all OA neurons in the brain, we previously identified three OA cell types - two unpaired and one paired - that directly innervate the mushroom bodies (Selcho et al., 2014). sVUM_mx1_ and sVUM_md1_ (also called OAN-a1 and OAN-a2) innervate the calyces, while sVPM_mx_ (also called OAN-g1) projects to the base of the vertical lobe. These three cell types were later confirmed via EM connectome data, which additionally revealed a fourth OA (paired) neuron innervating the tip of the vertical lobe (OAN-e1; (Eichler et al., 2017)). We therefore investigated the response dynamics of larval mushroom body Kenyon cells to OA and TA, as all OA-positive neurons in the subesophageal clusters are also TA-positive (Selcho et al., 2014). To this end, we genetically expressed the Ca^2+^ sensor GCaMP6m in Kenyon cells (*H24>GCaMP6m*) and bath applied OA and TA at various concentrations on isolated brains. DA was applied as a reference stimulus. Consistent with findings in the adult mushroom body (Tomchik and Davis, 2009), we observed only mild, concentration-dependent effects in intracellular Ca^2+^ levels in response to OA, TA and DA **(Fig.2A-F)**. Changes in fluorescence intensity were detected for all stimuli at a concentration of 10^-1^M, while TA also elicited changes at 10^-2^M **(Fig.2C-H)**. These results confirm that OA and TA signals modulate the activity of larval mushroom body Kenyon cells.

**Figure 2:**
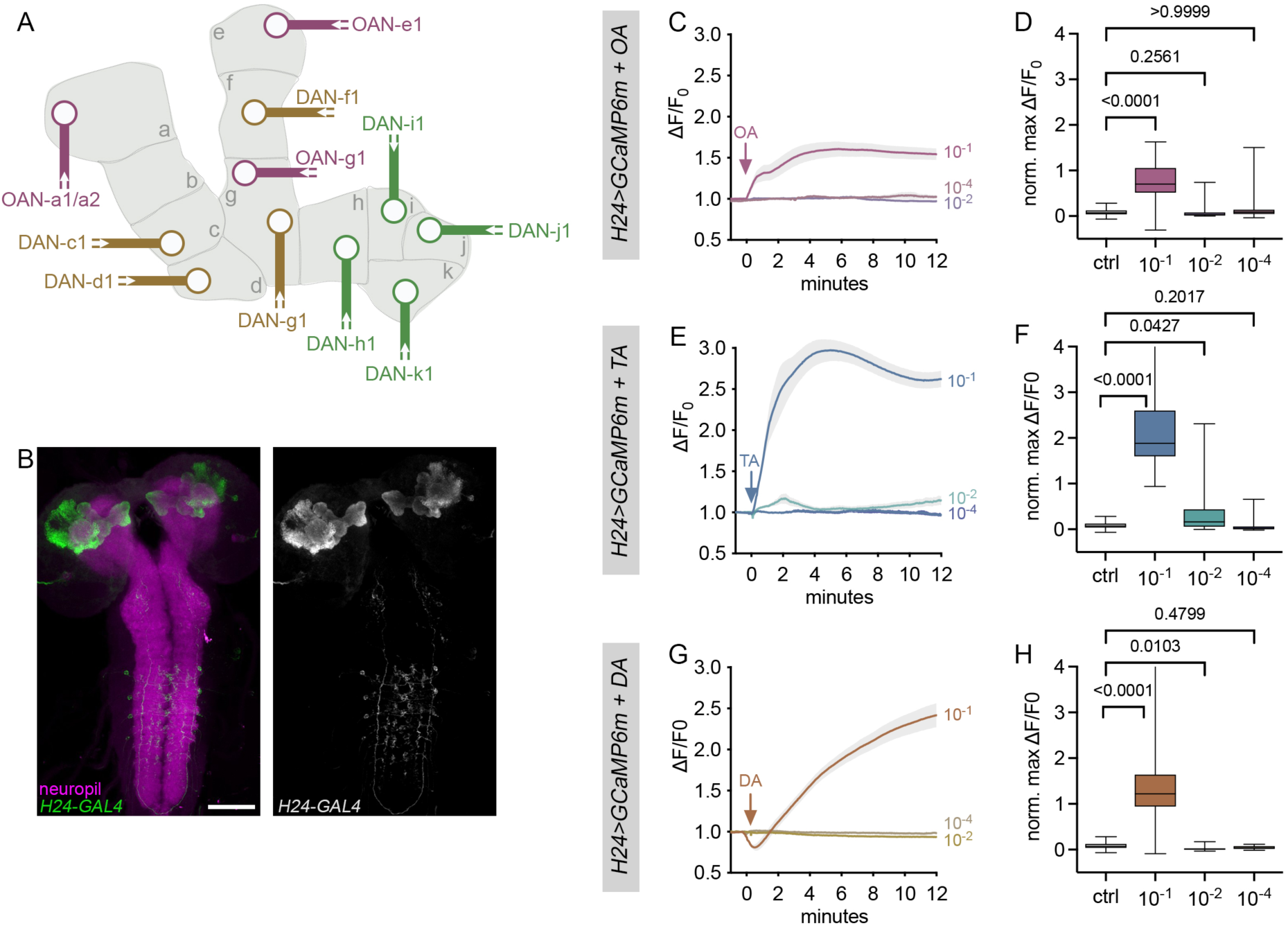
Concentration-dependent Ca^2+^ responses of larval mushroom body Kenyon cells to biogenic amines. (A) Schematic illustration of dopaminergic and octopaminergic mushroom body input neurons. Different compartments are innervated by different DA neurons (green: cells of the pPAM cluster; brown: cells of the DL1 cluster) and OA neurons (magenta), respectively. (B) Innervation pattern of H24-GAL4-positive neurons (green: H24>10xmyr::GFP; magenta: anti-synapsin / anti-ChAT / anti-FasII). H24-GAL4 labels all Kenyon cells of the larval mushroom body and few additional cells in the ventral nerve cord. (C) Changes in fluorescence intensities of Kenyon cells (regions of interest were set in the calyx) are visible to bath application of 10^-1^M OA, while no responses are detectable at 10^-2^M and 10^-4^M OA. (C,D) Fluorescence intensities change upon bath application of 10^-1^mM and 10^-2^M TA, respectively, while no response was detectable to bath application of 10^-4^M. (E,F) Changes in fluorescence intensities were detectable to bath application of 10^-1^M DA, while lower concentrations did not induce specific responses. Scale bar: 50µm.

The conundrum remains: how does activation of OA neurons induce an appetitive memory despite the absence of direct innervation of the medial lobes? To address this, we used available split-GAL4 lines specific to OAN-a1,a2 and OAN-g1 (no line was available for OAN-e1) (Eschbach et al., 2020) and activated those OA neurons that directly innervate the mushroom body Kenyon cells during substitution learning. Consistent with our previous results *SS24765>chop^XXL^*and *SS25844>chop^XXL^* larvae showed aversive memory expression compared to genetic controls. Specifically, activation of OAN-a1,a2 and OAN-g1 each elicited aversive memory expression during CS2 learning **(Fig.3G,H)** comparable to *Tdc2>chop^XXL^* larvae **(Fig.1F)**. However, in contrast to *Tdc2>chop^XXL^*, no appetitive memory expression was observed during CS1 learning **(Fig.3H)**. The absence of appetitive memory formation in CS1 learning aligns with the lack of medial lobe innervation by OA neurons, further reinforcing the notion that OA-mediated appetitive memory formation is anatomically constrained.

**Figure 3:**
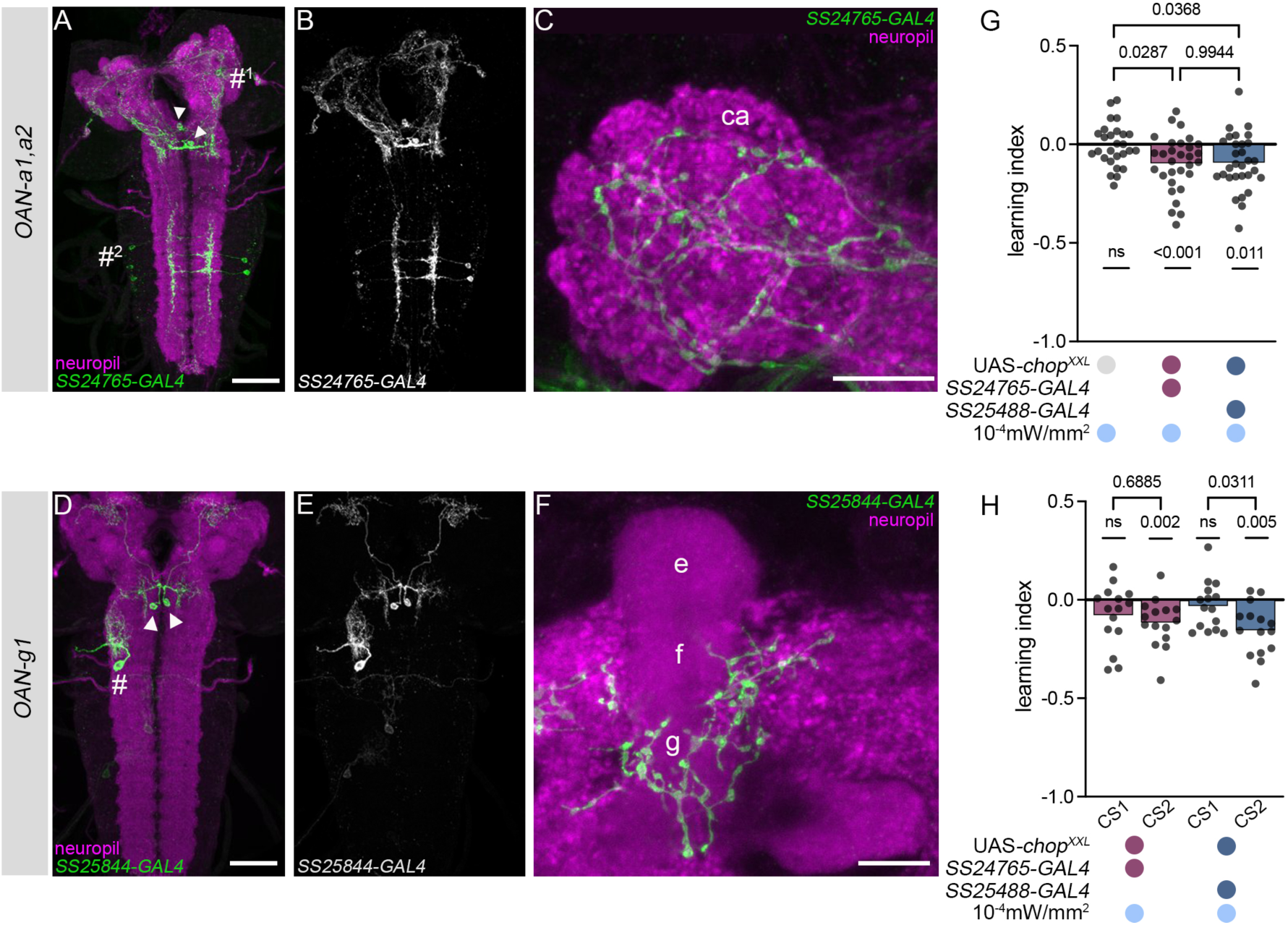
Optogenetic activation of specific OA neurons induces only aversive memory expression. (A-C) Innervation pattern of SS24765-GAL4-positive cells (green: SS24765>10xmyr::GFP; magenta: FasII/ChAT neuropil staining). The split-GAL4 line labels OAN-a1 and OAN-a2 (sVUMmx1 and sVUMmd1; arrowheads) and a few additional cells in the dorsal protocerebrum (#^1^) and ventral nerve cord (#^2^). Innervation of the mushroom body calyx. (D-F) Innervation pattern of SS25844-GAL4-positive cells (green: SS25844>10xmyr::GFP; magenta: FasII/ChAT neuropil staining). The split-GAL4 line labels the paired sVPMmx neurons (arrowheads) innervating the vertical lobe (OAN-g1) and additional cells in the ventral nerve cord (#). (G) Optogenetic activation of OAN-a1 and OAN-a2 (SS24765>chop^XXL^) elicited significant aversive memory expression. (H) Optogenetic activation of OAN-a1 and OAN-a2 induced aversive memory expression in CS2 learning, but no significant memory expression during CS1 learning. (B) Similar effects were obtained by artificial activation of the OAN-g1 cell type. Scale bars: 50µm (A,D), 10µm (C,F)

We next pursued an alternative hypothesis regarding how OA neurons contribute to larval reward learning. Specifically, we considered the possibility that OA neuron activation indirectly facilitates reward learning by stimulating DA neurons of the pPAM cluster, which are known to provide teaching signals to the medial lobes during memory formation, as previously shown in flies (Burke et al., 2012; Meschi et al., 2024). To test this, we expressed GCamp6m in pPAM neurons (*R58e02>GCaMP6m*) and performed bath applications of OA and TA, respectively, on isolated brains. Both OA and TA elicited calcium transients in pPAM neurons of *R58e02-GAL4*, suggesting excitatory effects **(Fig.4A-B)**. However, closer evaluation of individual calcium traces revealed variability: some neurons showed little to no response, while others exhibited Ca^2+^ peaks at different time points, indicating heterogeneous response dynamics within the pPAM cluster **(Fig.S2)**. To achieve greater specificity, we next expressed GCaMP6m in split-GAL4 lines targeting individual DA neurons of the pPAM cluster. DAN-h1 (*SS01696>GCaMP6m;* **Fig.4C-D**) and DAN-j1 (*MB316B>GCaMP6m*; **Fig.4G-H**) exhibited significant increases in intracellular Ca^2+^ in response to OA, whereas TA elicited either only mild or no responses. In contrast, DAN-k1 (*SS01757>GCaMP6m*; **Fig.4I-J**) responded significantly to both OA and TA, while DAN-i1 did not exhibit any specific response (*SS00864>GCaMP6m*; **Fig.4E-F**). Our behavioural data further support the idea that activation of Tdc2-positive neurons is sufficient to induce aversive memory formation **(Fig.1F-G)**. To examine whether OA and/or TA influence aversive teaching signals, we expressed GCaMP6m in DAN-f1 and DAN-g1 neurons of the DL1 cluster (*MB054b>GCaMP6m*) and applied OA and TA on isolated brains. Previous studies have shown that DAN-f1 and DAN-g1 activation is both necessary and sufficient to establish taste punishment learning in the larva (Eschbach et al., 2020; Thoener et al., 2022; Weber et al., 2023b; Weiglein et al., 2021). Interestingly, bath applications of both OA and TA led to a significant reduction of Ca^2+^ levels in these neurons **(Fig.4K-L)**. These results may suggest that OA neurons / TA neurons do not function as aversive teaching neurons themselves but rather modulate DA neurons that encode aversive signals. Consistent with this, we found that interfering with Tdc2-positive neurons does not affect aversive learning. Neither conditional silencing via shibire^ts^ expression, nor constitutive hyperpolarization via Kir2.1, nor inhibition of synaptic transmission via tetanus toxin disrupted larval aversive odour-salt memory **(Fig.S3)**.

**Figure 4:**
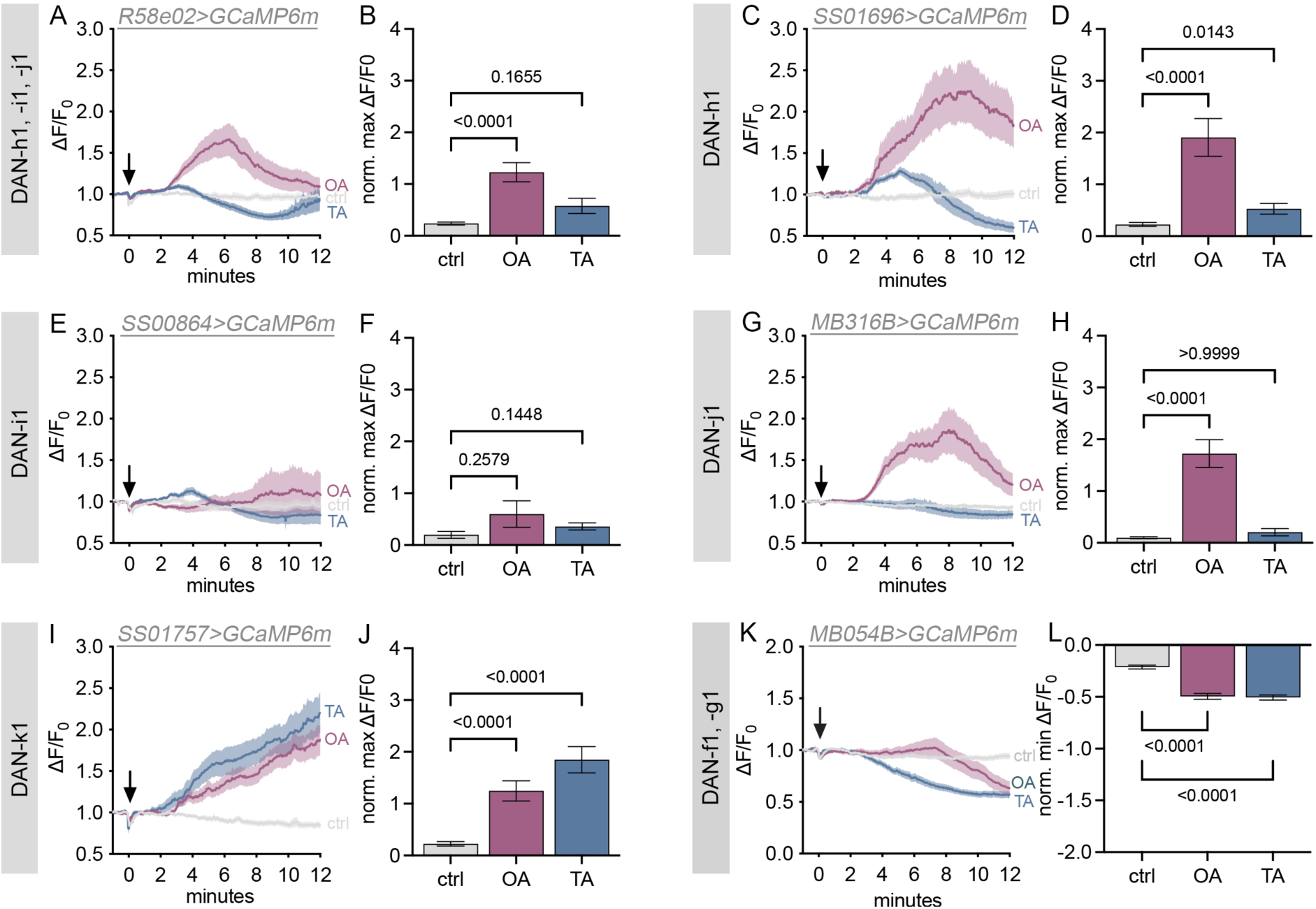
DA neurons are selectively modulated by OA and TA. (A-B) R58e02>GCaMP6m specimen were bath applied with OA and TA, respectively. Fluorescence intensities indicate that intracellular Ca^2+^ levels change upon application of OA, but not TA. To obtain cellular specificity, we used specific split-GAL4 lines for individual DA neurons. OA and TA application significantly altered fluorescence intensities of DAN-h1 (C-D) and DAN-k1 (I-J), while no fluorescence intensity changes were detectable in DAN-i1 (E-F). A specific change in response to OA only was observed in DAN-j1 (G-H). OA and TA bath application elicite a significant reduction in fluorescence intensities in DAN-f1 and DAN-g1 (K-L).

Our findings suggest that OA and TA signals modulate the activity of individual pPAM neurons (**Fig.4A-J**). To confirm that OA to pPAM signalling contributes to appetitive learning, we constitutively knocked down individual OA and TA receptors in pPAM neurons via RNA interference expression using *R58e02-GAL4* (targeting DAN-h1, DAN-i1, DAN-j1). For this screen, we primarily employed a one-cycle training regime, previously shown to yield robust immediate memory scores (Schleyer et al., 2011; Widmann et al., 2016). To systematically assess receptor function, we tested the effects of knocking down receptor levels on both odour-fructose and odour-arabinose associative learning. Interestingly, decreasing both TA receptor (TyrR and TyrRII) and several OA receptor (OAMB, Octɑ2R, Octß1R and Octß2R) levels did not impair memory performance in either paradigm **(Fig.S4A-L)**. In contrast, the knockdown of Octß3R in pPAM neurons (*R58e02>Octß3R-RNAi*) significantly impaired both odour-fructose and odour-arabinose memory **(Fig.5A,B)**. Consistently, Trojan-Octß3R larvae also failed to form appetitive memory in both paradigms **(Fig.5C,D)** (Diao et al., 2015). To further validate these findings, we repeated the Octß3R knockdown experiments using a three-cycle training paradigm. Unlike single-cycle training, repeated training alters the composition of the established memory phases (Weber et al., 2023a; Widmann et al., 2016), thereby providing a more rigorous test for the role of Octß3R in larval reward learning. As before, interfering with Octß3R reduced memory scores for both odour-fructose and odour-arabinose learning compared to genetic controls. However, memory performance remained above chance level, suggesting that Octß3R is specifically required for non-consolidated forms of memory **(Fig.5E-H)**. Importantly, both *R58e02>Octß3R-RNAi* and Trojan-Octß3R larvae exhibited normal odour and sugar preferences **(Fig.5I-P)**, ruling out general sensory or motivational deficits. Moreover, knockdown of Octß3R in mushroom body Kenyon cells did not impair memory, reinforcing the conclusion that its role is specific to pPAM neurons **(Fig.S4M-P)**. Together, these results support a specific role for Octß3R in pPAM neurons during larval appetitive learning and memory.

**Figure 5:**
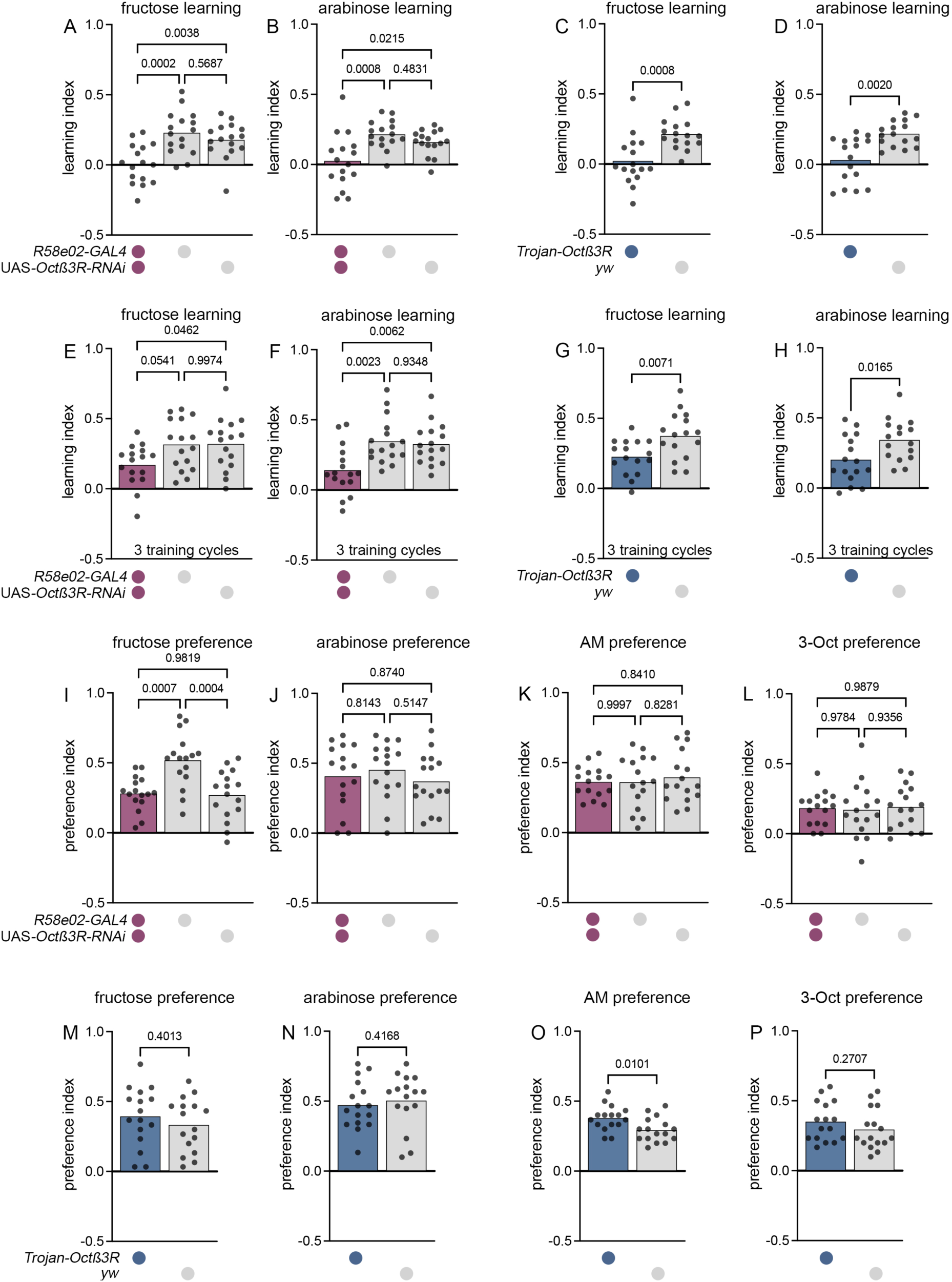
Octß3R is required in PAM neurons for appetitive odour-taste learning. (A) Downregulation of Octß3R in pPAM neurons (R58e02>Octß3R-RNAi) impairs odour-fructose and odour-arabinose learning. (B) In line with the specific knockdown of Octß3R in DA neurons, Trojan-Octß3R larvae are significantly reduced in odour-fructose and odour-arabinose learning compared to genetic controls. (E-H) Interfering with Octß3R signalling results in reduced learning scores after three cycles of training. Both R58e02>Octß3R-RNAi (I-L) and Trojan-Octß3R (M-P) larvae showed normal sensory preferences for odours and tastants.

As only DAN-h1 has previously been shown to be necessary for odour-sugar learning (Saumweber et al., 2018), we focused our analysis to this neuron type using *SS01696-GAL4* and determined whether OA modulation of DAN-h1 is required for appetitive memory performance. Interestingly, *SS01696>Octß3R-RNAi* larvae showed significantly reduced memory scores in odour-arabinose learning, while odour-fructose learning remained indistinguishable from controls **(Fig.6A,B)**. The specific impairment in arabinose-associated learning, along with the preserved fructose learning are consistent with previous findings (Selcho et al., 2014). However, these results appear somewhat inconsistent with the Octß3R knockdown effect using *R58e02-GAL4* **(Fig.5G-H)**. To further confirm the central role of DAN-h1 in OA-dependent modulation of appetitive memory, we performed Ca^2+^ imaging with simultaneous application of OA and downregulation of Octß3R in DAN-h1 (*SS01696>GCaMP6m;Octß3R-RNAi*). As expected, OA application failed to elicit Ca^2+^ transients under these conditions, underlining the necessity of this particular OA receptor for modulating DAN-h1 activity (**Fig.6C,D**). Taken together, our findings suggest that OA neurons exert a specific and selective modulatory role on distinct DA neuron types. In this context, DAN-f1 and DAN-g1 are essential for aversive learning, while DAN-h1 is crucial for appetitive learning.

**Figure 6:**
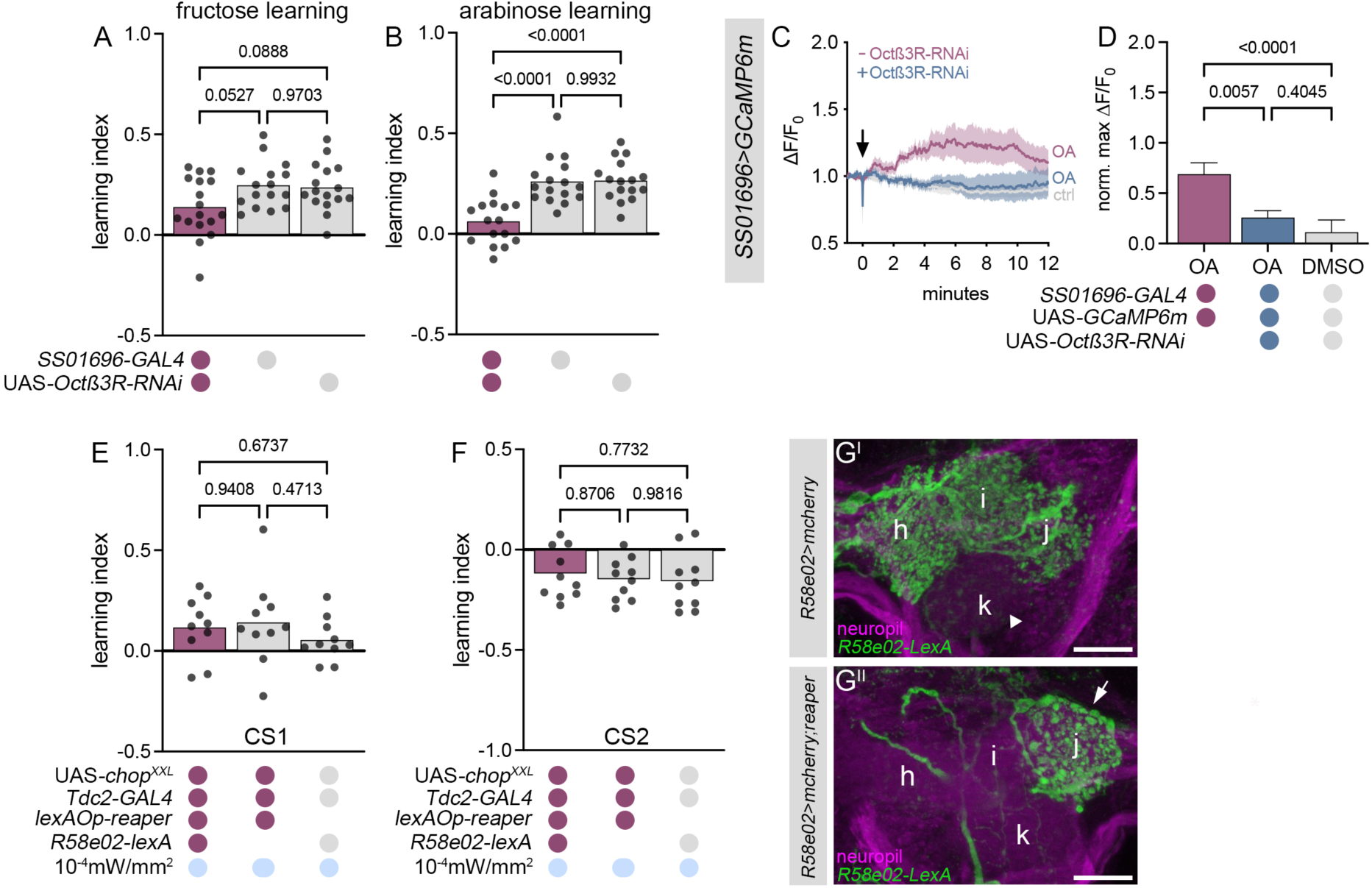
DAN-h1 is necessary for larval odour-reward memories, while DAN-j1 is sufficient. (A,B) Specific downregulation of Octß3R in DAN-h1 (SS01696>Octß3R-RNAi) significantly reduced memory performance for arabinose learning. (C,D) Downregulation of Octß3R in DAN-h1 prevents changes in fluorescence intensities in SS01696>GCaMP6m specimens to OA bath application using Ca^2+^ imaging. (E,F) Optogenetic activation of Tdc2-positive neurons and simultaneous ablation of R58e02-positive neurons did not alter CS1 appetitive memory or CS2 aversive memory expression. (G) Immunohistochemical analysis revealed that ablation of R58e02-positive neurons (G^I^; i.e. DAN-h1, DAN-j1, DAN-i1) was incomplete. Expression of lexAop-reaper induced apoptosis in DAN-h1 and DAN-i1 while DAN-j1 escaped ablation (G^II^, indicated by the arrow). Arrowhead: lack of innervation of the k-compartment, as DAN-k1 is not included in the R58e02-LexA line. Scale bars: 10µm.

To further validate that OA:DAN-h1 signalling is fundamental for the induction of positive CS1 memory in our optogenetic substitution experiment, we performed a combined gain-of-function and loss-of-function approach. Specifically, OA neurons were optogenetically activated while pPAM neurons, including DAN-h1, were simultaneously ablated. Contrary to our expectations, *Tdc2>chop^XXL^; R58e02>reaper* larvae still showed appetitive memory expression for CS1 learning (and as expected, an aversive memory expression for CS2 learning; **Fig.6E,F**). Surprisingly, immunohistochemical analysis revealed that DAN-j1 escaped *reaper*-induced apoptosis, while DAN-h1 and DAN-i1 were ablated as expected (**Fig.6G**). These results suggest that, in addition to DAN-h1, which is both necessary and sufficient for appetitive learning **(Fig.6A-C)** (Saumweber et al., 2018), DAN-j1 is also sufficient to orchestrate OA-dependent appetitive memory. Notably, DAN-k1, which is not covered by *R58e02-GAL4* line and therefore not targeted in either the receptor knockdown nor ablation experiments, was found to be dispensable for appetitive memory expression **(Fig.S5)**.

## DISCUSSION

In this study, we confirm the modulatory role of OA in learning and memory in *Drosophila* larvae, consistent with previous research. Importantly, we refine the current understanding of OA’s mode of action by identifying the functional mechanisms of OA neurons in larval aversive memory. We also provide novel, single cell level evidence for the involvement of the Octß3 receptor in modulating the dopaminergic pPAM cluster during appetitive memory formation. Our findings reveal for the first time the bimodal functionality of the OA system in both appetitive and aversive learning in larvae, highlighting its distinct sites of action.

The function of OA neurons in learning and memory was initially described by the mediation hypothesis (Claßen and Scholz, 2018; Sombati and Hoyle, 1984; Widmann et al., 2018), which proposed that individual OA neurons primarily convey reward signals and function directly as teaching neurons (Hammer, 1993; Honjo and Furukubo-Tokunaga, 2009; Kim et al., 2013; Schwaerzel et al., 2003). However, accumulating evidence from both larval and adult *Drosophila* challenges this model. First, OA signalling has been implicated in both adult appetitive and aversive learning, as shown by studies on *TßH* mutants and *Octß2R* and *Octß1R* mutants (Iliadi et al., 2017; Sabandal et al., 2020; Selcho, 2024; Wu et al., 2013). Consistent with this, we now demonstrate that optogenetic activation of OA neurons, when paired with an odour stimulus, is sufficient to induce aversive memory in CS2 training paradigm **(Fig.1F,G)**. Second, interfering with OA signalling during adult odour-sucrose learning (Burke et al., 2012), or ablating OA neurons in larval odour-fructose (Selcho et al., 2014) or aversive odour-salt learning **(Fig.5D-F)**, does not impair memory formation, suggesting that OA neurons are not essential for these types of learning. Third, at the anatomical level, OA neurons do not directly innervate the medial lobes of the mushroom bodies in larvae, despite these regions being critical for appetitive memories storage (Selcho et al., 2014; Widmann et al., 2018).

Based on these findings, the role of OA neurons can be rather reinterpreted in light of the orchestration hypothesis (Claßen and Scholz, 2018; Hernandez et al., 2023; Sombati and Hoyle, 1984). According to this model, OA neurons do not directly mediate teaching signals but instead modulate other neurons of the mushroom body circuitry in a context-dependent manner. Our results now provide direct support for this hypothesis. Previous studies have provided extensive evidence that DA neurons of the pPAM cluster function as teaching neurons during memory formation, consistent with the mediation hypothesis (Burke et al., 2012; Ichinose et al., 2015; Rohwedder et al., 2016; Yamagata et al., 2015). In larvae, the pPAM cluster comprises four neurons (DAN-h1, DAN-i1, DAN-j1 and DAN-k1) which directly innervate the medial lobes of the mushroom bodies. The activity of these neurons during memory formation is both necessary and sufficient to shift mushroom body output toward attraction. Our study now provides further information on the functional division of the pPAM cluster. We show that individual pPAM neurons are selectively modulated by OA. First, DAN-h1 activity is enhanced by OA through Octß3R, and knockdown of Octß3R specifically in DAN-h1 significantly impairs appetitive memory formation **(Fig.6A-D)**. Further, DAN-h1 activity has shown to be both necessary and sufficient to induce larval attraction suggesting that DAN-h1 is the central cell of the pPAM cluster during memory formation (Saumweber et al., 2018). DAN-j1 appears to contribute to reward learning as well, as evidenced by its response to OA stimulation **(Fig.4G)** and the preservation of learning following ablation of DAN-h1 and DAN-i1 **(Fig.6E-G)**. We also confirm that DAN-k1 is not sufficient to induce memory expression (**Fig.S5A-B)**. However, we demonstrate that DAN-k1 is activated by OA and TA, potentially enabling context-dependent modulation of sensory signal evaluation. Notably, OA and TA both elicit an excitatory response in DAN-k1 (**Fig.4I**), a property unique to this neuron within the pPAM cluster and they do not act antagonistically in this context as they do, for example, in other processes such as locomotion (Kutsukake et al., 2000; Nagaya et al., 2002; Pauls et al., 2015; Schützler et al., 2019; Selcho et al., 2012; Selcho and Pauls, 2019). Strikingly, in larvae where only DAN-j1 and DAN-k1 are functional - and where DAN-k1 alone does not impact reward learning - OA neuron stimulation still induces a positive CS1 memory **(Fig. 6E-G)**, highlighting the importance of DAN-j1. In contrast, DAN-i1 does not respond to OA or TA, suggesting that it is not modulated by these catecholamines, despite previous evidence supporting its role in appetitive memory **(Fig.4E,F)**. This underscores a nuanced, cell-type-specific modulation within the pPAM cluster.

OA orchestrates pPAM activity along a gradient of modulation strength, following the sequence of DAN-h1, DAN-j1, DAN-k1, and DAN-i1, from strong to absent responsiveness. Given that pPAM neurons are synaptically segregated from each other, as revealed by EM reconstructions (Eichler et al., 2017), this organization likely enables individual, context-dependent modulation of teaching signals within distinct mushroom body compartments. This organisation differs markedly from that in adult *Drosophila*, where crosstalk between dopaminergic input neurons has been proposed and shown to be essential for the specificity of signals to the distinct medial lobe compartments (Cohn et al., 2015). Moreover, additional modulators likely contribute to tuning pPAM activity, potentially extending beyond the modulation of DAN-i1. For instance, recurrent feedback from Kenyon cells via sNPF signalling has been implicated in memory stabilization within the larval pPAM cluster (Lyutova et al., 2019). This is consistent with our observation that optogenetic activation of pPAM neurons at high light intensity resulted in learning scores exceeding those of both lower light stimulation and even wild-type sugar learning (**Fig.1**). These findings may imply that the dopaminergic system can dynamically adapt to modulatory input through both feed-forward (OA and TA) and feedback (sNPF) mechanisms, highlighting the integrative and dynamic nature of reinforcement signalling in the larval brain.

Can we rule out that OA neurons function according to the mediation hypothesis? For larval appetitive learning, a direct modulation of mushroom body Kenyon cells, such as that mediated by OAMB in flies (Kim et al., 2013), has not been documented, making this scenario unlikely. However, our findings provide the first evidence that such a mechanism may be possible for larval aversive memory. Specifically, optogenetic activation of OAN-a1, OAN-a2 (targeting the calyx), as well as OAN-g1 (targeting the lower vertical lobe), induces aversive CS2 memory **(Fig.3)**. Interestingly, the optogenetic activation of these neurons using Cs-Chrimson in a previous study (Eschbach et al., 2020) showed a similar trend toward aversive learning, but the effect was not significant. This highlights the selective effect of different optogenetic tools on OA neurons (**Fig.1**). Further experiments are needed to determine whether the effect results from direct OA signalling to mushroom body Kenyon cells or occurs indirectly via modulation of specific dopaminergic teaching neurons. In the calyx, modulation may occur via the GABAergic APL neuron, and in the lower vertical lobe via the dopaminergic DAN-g1 (Eichler et al., 2017; Mancini et al., 2023a; Saumweber et al., 2018). Connectome data support this, showing direct synaptic connections between these OA neurons and either APL or DAN-g1 (Eichler et al., 2017; Mancini et al., 2023b). In contrast, direct OA-mediated signalling to Kenyon cells remains unlikely in larvae, as there is currently no evidence for a larval counterpart to the Octß1R-dependent mechanism described in flies (Sabandal et al., 2020). Instead, our data show that DAN-g1 is modulated by OA signalling **(Fig.4K)**, supporting an indirect modulatory role of OA in aversive memory formation.

Previous studies have shown that the larval aversive learning system is similarly organised to its appetitive counterpart described here and that four DA neurons of the DL1 cluster provide aversive teaching signals (Eschbach et al., 2020; Thoener et al., 2022; Weiglein et al., 2021). Among these, the combined activity of DAN-f1 and DAN-g1 has been shown to be both necessary and sufficient for aversive odour-salt learning (Weber et al., 2023a). While these neurons act as central teaching neurons, DAN-c1 and DAN-d1 appear to act in a supportive manner, mirroring the hierarchical structure of the pPAM cluster, where DAN-h1 functions centrally and other neurons supportively **(Fig.7)**. Our data further reveal that DAN-f1 and DAN-g1 receive input from both OA and TA, but in contrast to cells of the PAM cluster, DAN-f1 and DAN-g1 are inhibited by OA and TA. This inhibition aligns with appetitive learning, where OA/TA signalling activates reward-related DA neurons and suppresses aversive DA neurons, thus promoting attraction **(Fig.7)**. Given that optogenetic activation of OAN-g1 induces aversive CS2 memory **(Fig.3D-H)**, it seems unlikely that this effect is mediated by DAN-g1. Instead, the effect may arise from direct synaptic modulation of mushroom body output neurons or Kenyon cells. Indeed, connectome data support this, as the strongest connection of OAN-g1 is to MBON-g1,g2, accounting for 28% of all output synapses, while connections to DAN-g1 (0.02% of output synapses) and mushroom body Kenyon cells (0.04% of output synapses) are rather limited (Eichler et al., 2017). In summary, these findings suggest that OA and TA orchestrate learning and memory by engaging distinct mechanisms at multiple circuit levels. Their effects depend on the valence of the learning experience, the specific neuronal targets, and are modulated by context, prior experience, and the organism’s internal state.

**Figure 7:**
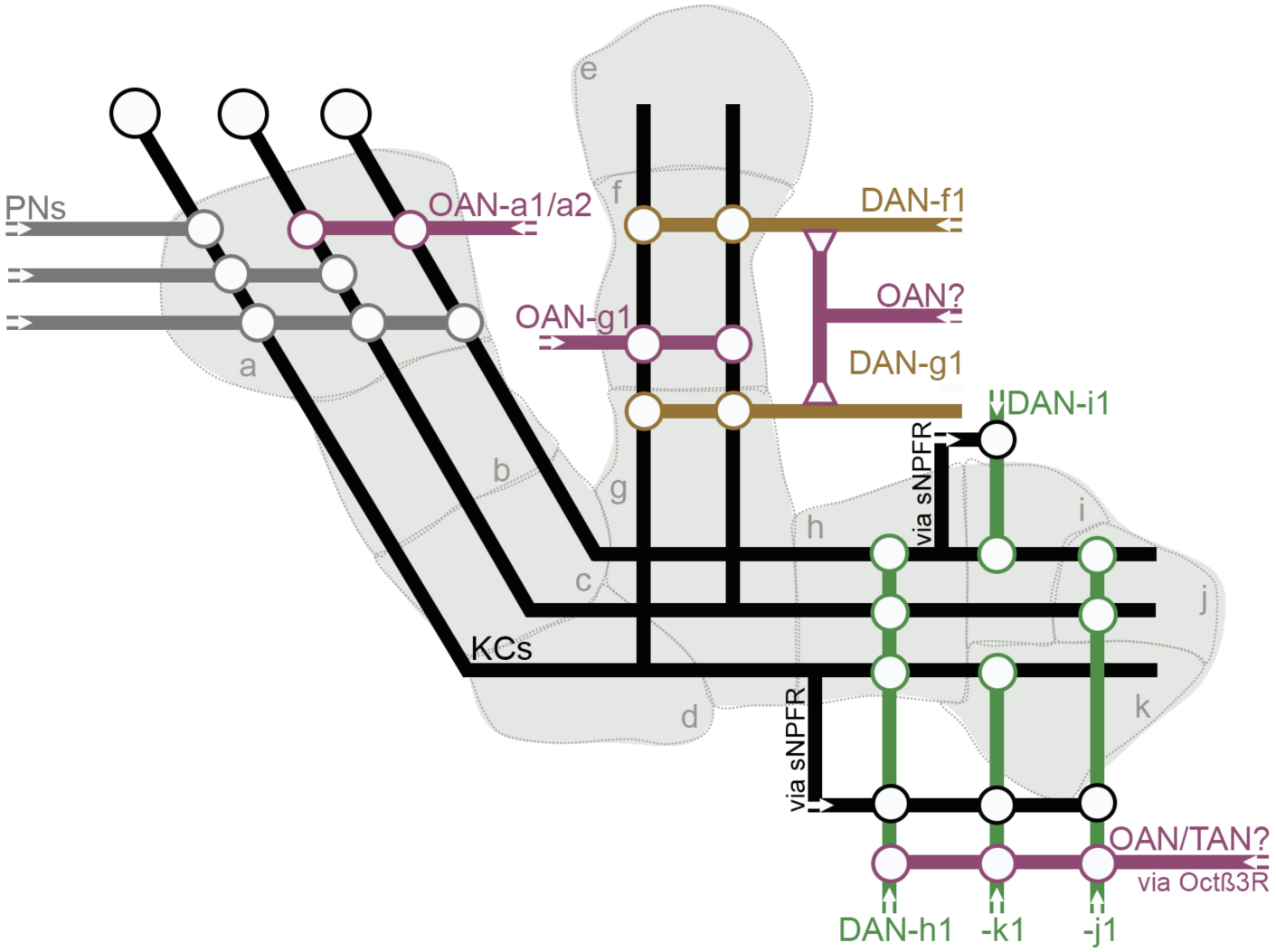
Simplified illustration of the connectivity within the mushroom body circuitry. While DA neurons of the PAM cluster (green; DAN-h1, -i1, -j1 and -k1) serve as teaching neurons during appetitive learning, DA neurons of the DL1 cluster (brown; DAN-f1/DAN-g1) serve as teaching neurons during aversive learning. OA neurons orchestrate appetitive olfactory learning by stimulation of the pPAM cluster DANs (through Octß3R) and suppression of DL1 cluster DANs. ca: calyx; DAN: dopaminergic neuron; KCs: Kenyon cells; OAN: octopaminergic neuron; PNs: projection neurons; sNPFR: short neuropeptide F receptor; TAN: tyraminergic neuron

The question remains: why is the activation of OA neurons sufficient to induce conflicting memories **(Fig.1)**? Notably, natural stimuli can also induce conflicting memories in larvae, even when applied using identical training protocols. For example, a mixture of 20 amino acids has been shown to elicit both appetitive and aversive memories, depending on the context (Toshima et al., 2019). Similarly, in adult *Drosophila*, a single alcohol conditioning protocol initially generates an aversive memory, which later transitions into a long-term appetitive memory (Kaun et al., 2011). A comparable phenomenon is observed with sugar-DEET mixtures, which establish simultaneous positive and negative memories in distinct compartments of the mushroom body (Das et al., 2014). These findings suggest that DA neurons can encode multiple memories in parallel, targeting different mushroom body compartments, each characterized by unique valence, dynamics, storage capacities, and flexibility (Aso and Rubin, 2016). A particularly striking result from the amino acid study is the context-dependent retrieval of these memories: positive memory is retrieved only in the absence of the rewarding stimulus, while negative memory is recalled in the presence of an aversive cue (Toshima et al., 2019). Additional studies in larvae have extended this principle to a variety of stimuli, including sugars, salts, and bitter compounds (Pauls et al., 2010a; Schleyer et al., 2015, 2011), and even the optogenetic activation of neurons (Lyutova et al., 2019), showing that memory retrieval is tightly gated by context and internal state. However, aversive odour-electric shock memories remain robustly expressed even in the absence of the aversive US during testing (Aceves-Piña and Quinn, 1979; Pauls et al., 2010a). This also applies to the optogenetic activation of OA neurons, where CS2 training induces aversive memory even on a neutral test plate **(Fig.1)**. In summary, numerous studies in *Drosophila* demonstrate that identical teaching stimuli can produce opposing memories, which are selectively retrieved in a context-dependent manner. Our results therefore suggest that OA may thus play a key role in orchestrating both the formation and retrieval of divergent memories, ensuring behaviour remains adaptively tuned to context.

On top, our findings confirm that a single teaching stimulus can induce opposing memories depending on the stimulus presentation regime: CS1 learning elicits appetitive memory expression, whereas CS2 learning elicits aversive memory expression, and not - for example - vice versa. The sequence effect appears to be specific to the activation of OA neurons, as the effect does not occur with optogenetic activation of pPAM neurons (**Fig.1)** or mushroom body Kenyon cells (Lyutova et al., 2019). A recent study on OA-and DA-dependent conditioning in crickets suggests a circuit model for the parallel formation of appetitive and aversive memories and their competition for the conditioned response. Here, different types of OA and DA neurons impact on both the formation and retrieval of memories through mutual activation and inhibition within the mushroom body circuitry (Rahman et al., 2024). This may well be consistent with the findings of this study on the selective modulation of extrinsic dopaminergic neurons by OA and TA. Alternatively, it has been shown that a temporal shift of less than a second is sufficient to switch behavioural responses from avoidance to attraction in optogenetic association learning experiment in adults. This bidirectional tuning is based on plasticity at Kenyon cell to mushroom body output synapses, triggered by activity of distinct DA neurons (Cohn et al., 2015). Optogenetic activation of OANs using *chop^XXM^* provides evidence that the sequential memory effect observed in our study (CS1 vs. CS2) is not due to the prolonged open state or to an exceptionally high expression of *chop^XXL^*. However, it is still tempting to speculate that the expression of opposing memories relies on temporal dynamics within the mushroom body circuitry at the level of mushroom body output neurons, which can also be directly triggered by optogenetic activation of OANs (**Fig.4**) (Ehmann and Pauls, 2020).

Another explanation may be that, in addition to the odour paired with the unconditional stimulus (CS+), a memory with opposite valence is also established for the unpaired odour (CS-), as it has been shown in adult *Drosophila* (Jacob and Waddell, 2020). Finally, in adult *Drosophila*, DANs are known to encode relative reinforcement signals (i.e., whether an experience is better or worse in comparison to another). Thus, whether an odour is initially presented in a neutral context or in conjunction with OA neuron modulation could significantly influence the resulting memory formation (Villar et al., 2022).

In vertebrates, it has been shown that vertebrate reward-mediating neurons are also sensitive to aversive input, a concept described as the “single-dimension” model. In contrast, the “two-dimension” model posits that such neurons are exclusively tuned to reward processing and thus other neurons must encode for aversive input (Fiorillo, 2013). Our findings now support the ‘two-dimension’ model at least for appetitive memories, in which the OA neurons add another level of complexity to the mushroom body circuitry through different modes of modulation (Marder, 2012). DA neurons of the pPAM cluster encode for reward, while neurons of the DL1 cluster encode for punishment. However, OA neurons act upstream of these neurons and selectively modulate their teaching signal. Our Ca^2+^ imaging data suggest that OA activates three of the four pPAM neurons, while inhibiting DL1 cluster DA neurons. This shifts the balance at the mushroom body input level in favour of approach and approach memory. Further research is needed to understand whether OA orchestrates also at the level of output neurons. This could explain the systemic effect of OA on learning and memory, where the complexity arising from multiple interactions partially obscures a precise characterisation of its cellular mechanism of action.

## MATERIAL AND METHODS

### Fly strains

Flies were cultured on standard food at 25°C and 60% humidity in a standard light:dark cycle. Experimental transgenic lines were obtained by crossing driver lines to UAS lines, whereas genetic controls were obtained by crossing those lines to *w^1118^*.

**Table 1:** Transgenic and mutant lines used in this study.

| Driver lines | Genotype | Reference |
| --- | --- | --- |
| 201y-GAL4, UAS-mCD8::GFP | y[1] w[67c23]; P{w[+mC]=UAS-mCD8::GFP.L}LL5<br>P{w[+mW.hs]=GawB}Tab2[201Y] | BDSC 64296 |
| H24-GAL4 | P{w[+mW.hs]=GawB}H24 | BDSC 51632 |
| MB316B-GAL4 | w[1118]; P{y[+t7.7] w[+mC]=R58E02-p65.AD}attP40/CyO; P{y[+t7.7] w[+mC]=R93G08-GAL4.DBD}attP2 | BDSC 68317 |
| *MB054B-GAL4* | Intersection strains: R76F05_R15B01 | (Eschbach et al., 2020; Pfeiffer et al., 2010; Weber et al., 2023a) |
| *R58e02-GAL4* | w[1118]; P{y[+t7.7] w[+mC]=GMR58E02-GAL4}attP2 | BDSC 41347 |
| *R58e02-LexA* | w[1118]; P{y[+t7.7] w[+mC]=GMR58E02-lexA}attP40 | BDSC 52740 |
| *R49D11-GAL4* | w[1118]; P{y[+t7.7] w[+mC]=GMR49D11-GAL4}attP2 | BDSC 38684 |
| *SS00864-GAL4* | Intersection strains: 13D05_36B06 | (Eschbach et al., 2020; Pfeiffer et al., 2010; Saumweber et al., 2018; Truman et al., 2023; Weber et al., 2023a) |
| *SS01696-GAL4* | Intersection strains: 76F05_95H02 |  |
| *SS01757-GAL4* | Intersection strains: 48F09_27A11 |  |
| *SS24765-GAL4* | Intersection strains: 105H06_128G03 |  |
| *SS25844-GAL4* | Intersection strains: 040569_061921 |  |
| *Tdc2-GAL4* | w[*]; P{w[+mC]=Tdc2-GAL4.C}2 | BDSC 9313 |
| **Reporter lines** | **Genotype** | **Reference** |
| 10xUAS-IVS-myr::GFP | w[*]; P{y[+t7.7] w[+mC]=10XUAS-IVS-myr::GFP}attP2 | BDSC 32197 |
| UAS-*chop*<sup>XXL</sup> | w[1118]; PBac{yellow[+]-attP-9A Chop2_D156C = pRK010 w[+]/CyO w-GFP (w[-]) | BDSC 58374 |
| UAS-*chop*XXL;<br>lexAOp-reaper |  | (Lyutova et al., 2019) |
| lexAOp-*chop*<sup>XXL</sup> |  | (Selcho et al., 2017) |
| UAS-*chop*<sup>XXM</sup> | w[1118]; PBac{yellow[+]-attP-9A}VK00018 chop2[D156H (XXM)]= pTL538 w[+]/CyO w-GFP (w[-]) | (Dawydow et al., 2014) |
| UAS-GCaMP6m | w[1118]; P{y[+t7.7] w[+mC]=20XUAS-IVS-GCaMP6m}attP40 | BDSC 42748 |
| UAS-*hid,rpr* | y, w1118, P{UAS-hid}, P{UAS-rpr} | (Pauls et al., 2015; Zhou et al., 1997) |
| UAS-*Kir2.1* | w*; P{UAS-Hsap\KCNJ2. EGFP}1 | (Baines et al., 2001) |
| LexAOp-*mcherry* | y[1] w[*]; wg[Sp-1]/CyO, P{Wee-P.ph0}Bacc[Wee-P20]; P{y[+t7.7] w[+mC]=13XLexAop2-6XmCherry-HA}attP2 | BDSC 52271 |
| UAS-*OAMB-RNAi* | y[1] v[1]; P{y[+t7.7] v[+t1.8]=TRiP.JF01732}attP2 | BDSC 31233 |
| UAS-*Octa2R-RNAi* | y[1] v[1]; P{y[+t7.7] v[+t1.8]=TRiP.HMC03079}attP2 | BDSC 50678 |
| UAS-*Octβ1R-RNAi* | y[1] v[1]; P{y[+t7.7]<br>v[+t1.8]=TRiP.HMJ22156}attP40 | BDSC 58179 |
| UAS-*Octβ2R-RNAi* | y[1] sc[*] v[1] sev[21]; P{y[+t7.7]<br>v[+t1.8]=TRiP.HMS01151}attP2 | BDSC 34673 |
| UAS-*Octβ3R-RNAi* | y[1] v[1]; P{y[+t7.7]<br>v[+t1.8]=TRiP.HMJ23640}attP40 | BDSC 62283 |
| UAS-*shibire* (*shi<sup>ts1</sup>*) | w[*]; P{w[+mC]=UAS-shi[ts1].K}3 | BDSC 44222 |
| UAS-*TNTE* | w; P{UAS-TeTxLC.tnt}E2 | BDSC 28837 |
| UAS-*TyrR-RNAi* | y[1] sc[*] v[1] sev[21]; P{y[+t7.7]<br>v[+t1.8]=TRiP.HMC04811}attP2 | BDSC 57496 |
| UAS-*TyrRII-RNAi* | y[1] sc[*] v[1] sev[21]; P{y[+t7.7]<br>v[+t1.8]=TRiP.HMC05838}attP2 | BDSC 64964 |
| **Mutant lines** | **Genotype** | **Reference** |
| *Trojan-Octβ3R-GAL4* | y[1] w[*];; Mi{Trojan-GAL4.un}Octbeta3R[MI06217-TG4.un] | BDSC 76680 |
| **Control lines** | **Genotype** | **Reference** |
| CantonS |  | BDSC 64349 |
| w<sup>1118</sup> | w[1118] | BDSC 3605 |
| yw | y[1] w[1] | BDSC 1495 |

### Olfactory learning and memory

Learning experiments were performed in Petri dishes filled with a thin layer of either 1.5% or 2.5% agarose (CAS 9012-36-6, Roth). Depending on the experiment, a 1-cycle or 3-cycle two-odour reciprocal training regime was used (Schleyer et al., 2011; Weber et al., 2023a). 10µl of amyl acetate (AM; CAS 628-63-7, Merck), diluted 1:80 in paraffin oil (CAS 8012-95-1, Merck), or 10µl of undiluted 3-octanol (3-OCT; CAS 589-98-0, Merck) were used as conditional stimulus (CS) and loaded into Teflon containers with perforated lids. For odour-sugar learning, 2M fructose (CAS 57-48-7, Roth) or 2M arabinose (CAS 10323-20-3, Merck) were used as unconditioned stimulus. For odour-salt learning 1.5M sodium chloride (CAS 7647-14-5, Roth) was used. Aspartic acid (CAS 56-84-8, Merck) was used for experiments on test plates containing amino acids.

Substitution learning: fly cultures were wrapped in red foil before the experiment to protect them from blue light and thus the constant neuronal activation through chop^XXL^ / chop^XXM^. 250µM of *all-trans*-retinal (dissolved in ethanol; CAS 116-31-4, Merck) was applied to the standard food for experiments using chop^XXM^ (Dawydow et al., 2014; Scholz et al., 2017). During the experiment, we optogenetically substituted the unconditioned stimulus with constant blue light (475nm) of either ∼10^-1^mW/mm^2^ or ∼10^-4^mW/mm^2^ for experiments using chop^XXL^ and ∼10^-2^mW/mm^2^ for experiments using chop^XXM^.

During the test larvae were exposed to both odours (CS+ and CS-) in a binary choice test. After 5min larvae were counted on each side and a preference index was calculated by subtracting the number of larvae on one side from the number of larvae on the other side divided by the total number of larvae. The final learning index was then calculated from the mean value of two reciprocal experiments in which each of the two odours served as CS+ once. The final number of repetitions varied between the experiments. While we typically performed N=20 in optogenetic experiments (10x CS1, 10x CS), the standard N number in the other learning experiments was usually 15.

### Ca^2+^ imaging

Functional imaging was performed as described in (Lyutova et al., 2019; Selcho et al., 2017). To monitor intracellular Ca^2+^ levels in mushroom body Kenyon cells or dopaminergic neurons in response to bath application of OA and TA, respectively, GCaMP6m was expressed in cells of interest using the GAL4/UAS system (Brand and Perrimon, 1993). Larval brains were dissected and subsequently put in a Petri dish containing 405μl hemolymph-like HL3.1 saline solution. Before the experiment, brains were maintained for about 15-20min for settling. Images with ROIs selecting cell bodies (for dopaminergic neurons) or the calyx (for mushroom bodies) were recorded with an AXIO Examiner D1 microscope (Carl Zeiss AG, Germany) with a Zeiss W Plan-Apochromat 20×1.0 DIC (UV) VIS-IR objective and x20/1.0 water immersion objective and a Axiocam 506 camera (Carl Zeiss AG, Germany). Data is presented as fluorescence intensity (F) normalized to the baseline before stimulus application (F_0_). To analyse changes in fluorescence intensities in response to application of either octopamine (770-05-8, Merck), tyramine (CAS 60-19-5, Merck) or dopamine (CAS 62-31-7, Merck), we compared the maximum difference from baseline to vehicle control (i.e. HL3.1 containing 0.1% DMSO (Dimethyl sulfoxide; CAS 67-68-5, Merck)). Unless stated otherwise in the figures, a final concentration of 10^-2^M of OA and TA was used.

### Immunohistochemistry

Third instar larvae were dissected in ice cold 1x PBS. After fixation in 4% paraformaldehyde for 40 min and rinsing with 0.3% PBT (1x PBS with 0.3% TritonX-100; Merck) at room temperature, specimens were blocked over night at 4°C in 0.3% PBT with 5% normal goat serum (NGS).

Consecutive overnight incubation with primary and secondary antibodies **(Table 2)** in blocking solution was intermitted by 6×10 min washing in 0.3% PBT. Specimens were washed 5x 10 min in PBT and 1x 10 min in PBS and stored at 4 °C in Vectashield H-1000 (Vector laboratories) until they were mounted. Samples depicted in Figure 5 were scanned in confocal mode using an Olympus BX63F (EVIDENT Europe GmbH, Hamburg, DE) equipped with a UPLXAPO 60x, NA1.42 oil objective as part of an upright Abberior INFINITY LINE microscope (Abberior Instruments, Göttingen, DE). Image stacks of mushroom bodies were taken at a pixel size of 80 nm and a z-step size of 1 µm. Other samples were scanned at a confocal laser scanning microscope (ZEISS LSM800, ZEISS Microscopy). Overview image stacks of the brain lobes were taken at a pixel size of 0.83 µm and a z-step size of 1.5 µm using a LD LCI Plan-Apochromat 25x/0.8 Imm Korr DIC M27 objective, while detailed image stacks of mushroom bodies were taken at a pixel size of 100 nm and a z-step size of 1 µm using a 63x/1.4 NA oil DIC M27 objective. Image J [version 2.9.0/1.53t; (Schindelin et al., 2012)] was used to adjust brightness and contrast.

**Table 2:** Antibodies used in this study.

| Antibody | Manufacturer | Working dilution |
| --- | --- | --- |
| Mouse anti-ChAT | 4B1, DSHB | 1:50 |
| Mouse anti-Synapsin | 3C11, DSHB | 1:50 |
| Mouse anti-Fasciclin II | 1D4, DSHB | 1:50 |
| Mouse anti-Bruchpilot | Nc82, DSHB | 1:50 |
| Rabbit anti-GFP | A11112, life technologies | 1:500 |
| Rabbit anti-RFP | ABIN129578<br>Antibodies Online | 1:250 |
| Goat-anti-rabbit-AlexaFluor488 | A11034, life technologies | 1:500 |
| Goat-anti-mouse-STARred | Abberior | 1:100 |
| Goat-anti-rabbit-STARorange | Abberior | 1:100 |

### Statistics

Data were analysed using Graph Pad Prism 9. Data were analysed for normal distribution using the Shapiro-Wilk test. For normally distributed data, we used ANOVA followed by Tukey’s or Šidák’s multiple comparisons test. When comparing only two genotypes, we used either an unpaired t-test or the Mann-Whitney test. For data that were not normally distributed, we used the Kruskal-Wallis test followed by Dunn’s multiple comparisons test. To test against chance level, we performed a one-sample t-test.

## Acknowledgements

We thank Nicole Naumann, Barbara Goettgens, Johanna Pfitzenmaier, Astrid Rohwedder and Desiree von Alpen for technical support and the Bloomington Stock center and Vienna Drosophila Resource Center for fly strains. This work was supported by the Deutsche Forschungsgemeinschaft (430156010, 426503586, 447465118 to R.J.K., 426722269, 504082293, 531591887 to D.P., 420900883 to M.S., 441181781, 426722269, 432195391 to A.S.T) and by EU funds from the ESF Plus Program (Grant No. 100649752 to A.S.T.). The authors declare no competing interests. R.J.K., M.S., A.S.T. and D.P. conceived and designed the experiments. U.F., A.G., S.A., I.K., M.L., D.P. performed the experiments. U.F., A.G., S.A. M.L., M.S., R.J.K., A.S.T., and D.P. analysed the results. A.S.T. and D.P. wrote the manuscript. All authors provided comments and approved the manuscript.

## Supplement Material

**Figure S1:**
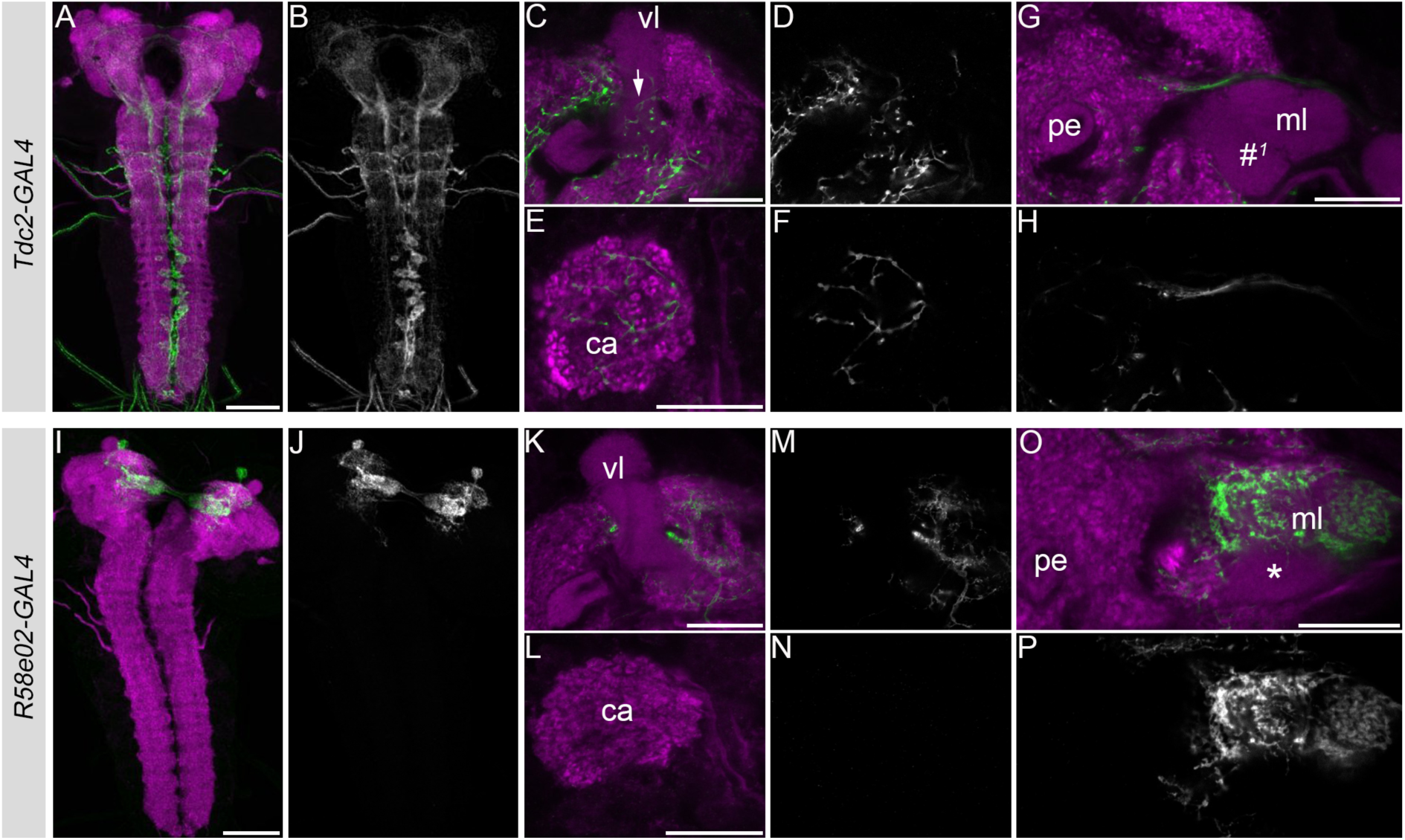
Innervation pattern of Tdc2-GAL4 and R58e02-GAL4. (A,B) Tdc2-GAL4 shows innervation in the brain and ventral nerve cord. (C-H) Selected regions of the larval mushroom bodies. (C,D) Tdc2-positive neurons innervate the vertical lobes in compartment g1 (indicated by the arrow) and the calyx (E,F). (G,H) No innervation is visible in the medial lobes of the mushroom bodies (indicated by #). (I,J) R58e02-GAL4 shows specific arborization in the larval mushroom bodies. (K-N) R58e02-positive neurons do not innervate the vertical lobe and calyx. (O,P) Innervation is visible in compartments of the medial lobes except for the k compartment (indicated by the asterisk). Both innervation patterns were described in detail before in (Rohwedder et al., 2016; Selcho et al., 2014, 2012). Scale bars = 50µm for whole mounts and 20µm for selections. Scale bar: 50µm or 10µm (C,E,G,K,L,O)

**Figure S2:**
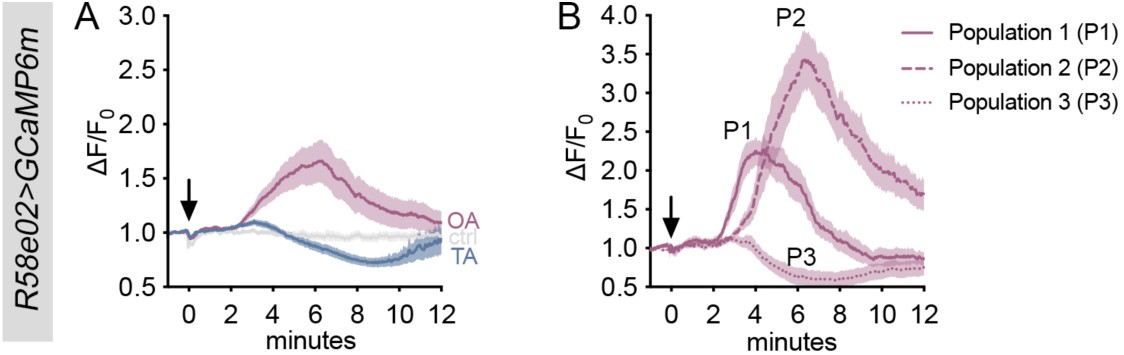
R58e02-labelled DANs show distinct response characteristics to OA. (A) Same data is presented in Figure 4A: R58e02>GCaMP6m specimen were bath applied with OA and TA, respectively. Fluorescence intensities indicate that intracellular Ca^2+^ levels change upon application of OA, but not TA. (B) Characteristic single traces of (A) indicating that the three cells included in R58e02-GAL4 (i.e. DAN-h1, DAN-i1 and DAN-j1) may have distinct response characteristics to OA.

**Figure S3:**
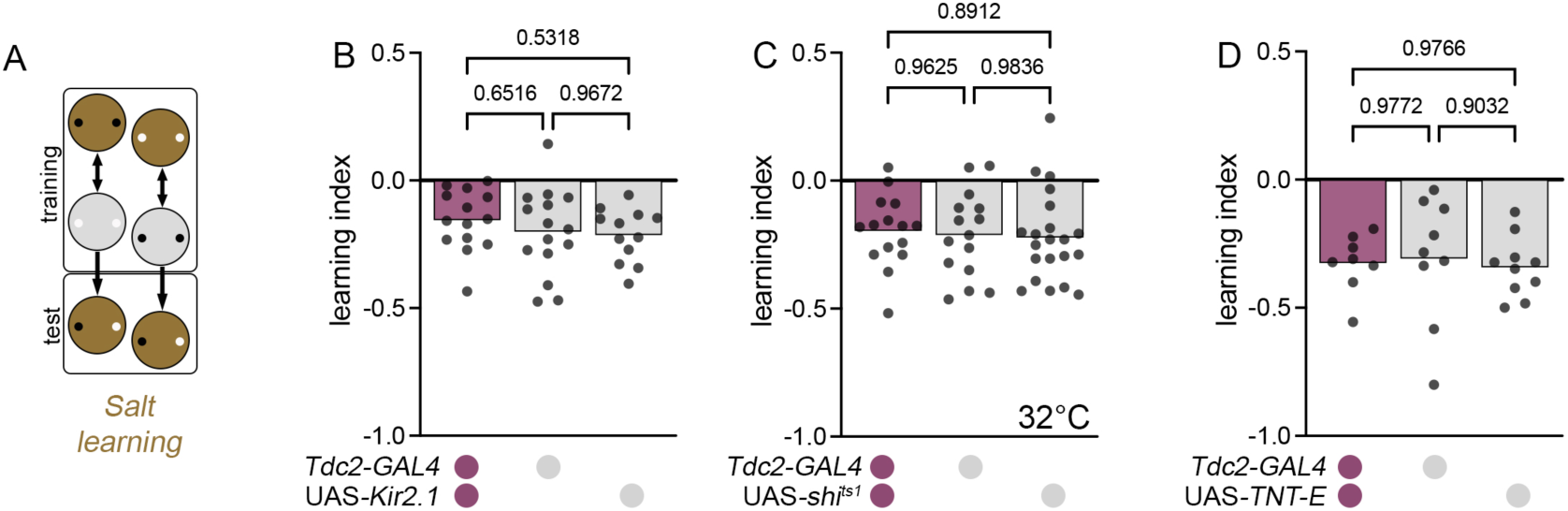
OA is not necessary for aversive odour-taste memories. (A) Schematic overview of the aversive learning protocol used in this study. Interfering with OA/TA signalling using Kir2.1 (B), Shibire^ts^ (C) and TNT-E (D) expression in Tdc2-positive neurons did not reduce aversive odour-salt learning compared to genetic controls.

**Figure S4:**
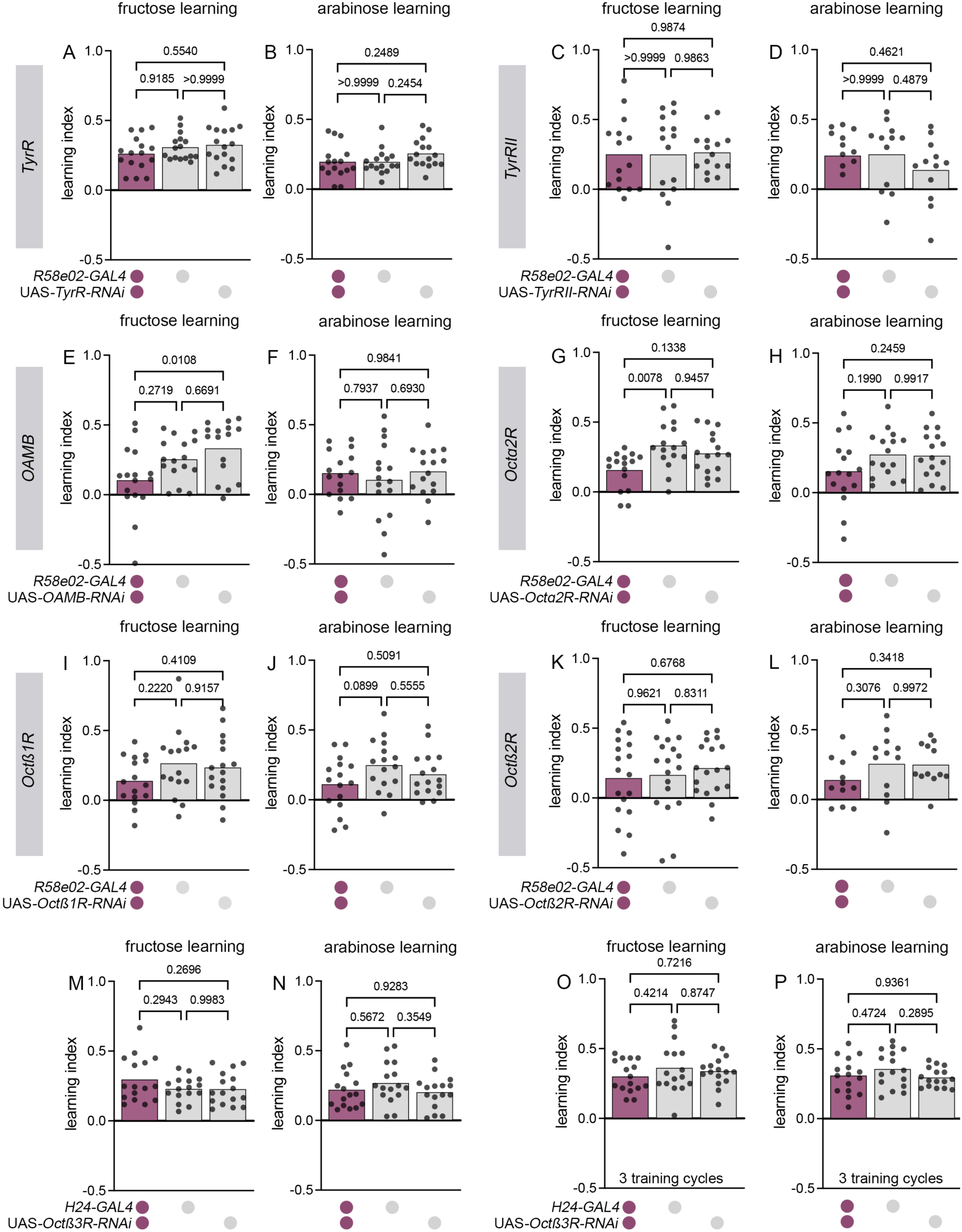
Only Octß3R is required in PAM neurons for appetitive odour-taste learning. (A-D) Downregulation of TA receptors TyR and TyRII does not impair odour-fructose and odour-arabinose learning. Similarly, specific knockdown of neither OAMB (E,F), Octɑ2R (G,H), Octß1R (I,J) nor Octß2R (K,L) affected appetitive memory performance. (M-P) Knockdown of Octß3R in mushroom body KCs did not affect appetitive memory performance using a one-cycle or three-cycle training regime.

**Figure S5:**
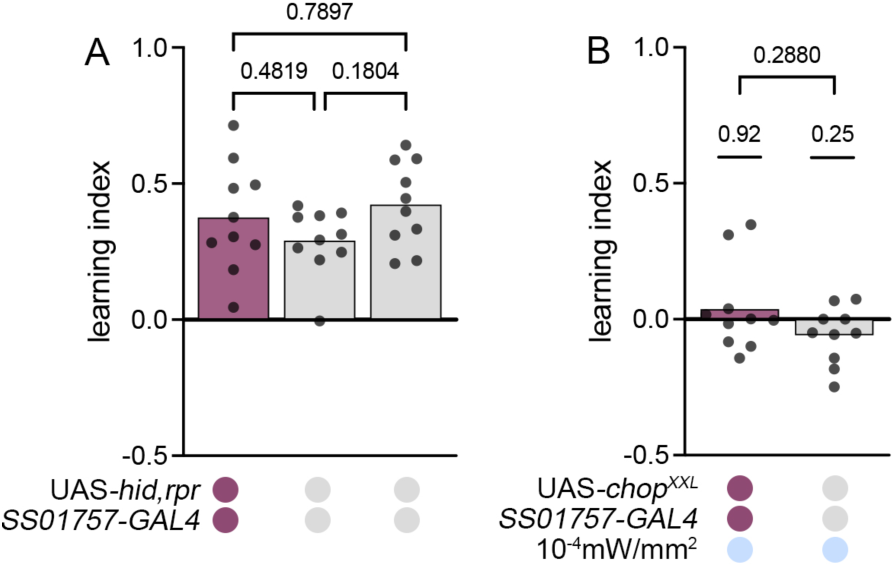
DAN-k1 is neither necessary nor sufficient for larval associative olfactory learning. (A) Ablation of DAN-k1 through the expression of ablation genes hid,rpr does not affect appetitive learning. (B) Optogenetic activation of DAN-k1 is not sufficient to induce memory expression in the larva.

